# Rapid body colouration change in *Oryzias celebensis* as a social signal for intraspecific competition

**DOI:** 10.1101/2023.12.23.573172

**Authors:** Ryutaro Ueda, Satoshi Ansai, Hideaki Takeuchi

**Affiliations:** Graduate School of Life Sciences, Tohoku University, 980-8577 Miyagi, Japan; Laboratory of Genome Editing Breeding, Graduate School of Agriculture, Kyoto University, Kyoto, 606-8507, Japan

## Abstract

In some species, such as chameleon and cichlid fish, rapid body colouration changes (within seconds or minutes) serve as visual social signals in male-male competition. This study investigated the relationship between aggressive behaviour and body colouration changes in *Oryzias celebensis*, an Indonesian medaka fish. We analysed aggressive behaviours and corresponding body colouration changes during attack events in a controlled laboratory setting using groups of 3 adult fish in a small tank. In a triadic relationship consisting of 2 males and a female, males with blackened markings attacked more frequently than males without blackened markings and females. Additionally, we observed that the males with blackened markings were seldom attacked by males without blackened markings and females. These tendencies persisted even in groups consisting of 3 males. Our results suggest that the blackened markings in male *O. celebensis* not only indicate the level of aggression but also serve as a social signal to suppress attacks by other individuals.

## Introduction

Across a wide range of taxa from vertebrates to invertebrates, animals such as fish, amphibians, reptiles, and cephalopods possess the ability to change their body colouration in response to external factors [1-4]. In most species, body colouration changes are primarily under hormonal control, but in some species of cephalopods, reptiles, and fish, neural systems directly control the chromatophore changes to respond very rapidly (within seconds). In chameleon males, rapid body colouration changes serve as social signals to indicate social status [2]. Chameleon males that are defeated in male-male competition rapidly change their colouration from bright yellow to brown [2]. Very few studies have investigated the neural mechanisms underlying rapid body colouration changes using molecular genetics.

In the present study, we used a medaka fish, *Oryzias celebensis*, endemic to southwest Sulawesi, Indonesia, as an experimental model. Medaka fish (family Adrianichthyidae) are widely distributed in East and Southeast Asia, and 20 of the 39 Adrianichthyidae species are endemic to Sulawesi [5, 6]. These fish exhibit significant diversification in sexually dimorphic traits such as morphology and body colouration, making them an excellent model for exploring the evolutionary genetic mechanisms underlying sexual dimorphism [7]. For one of these endemic species, *O. celebensis*, the reference genome assembly was generated in a previous study [8]. Interestingly, some male *O. celebensis* exhibit distinctive blackened markings on their fins and sides (figure 1a), and the colouration of these markings changes rapidly over a short period within a few seconds. Here we established a behavioural experimental paradigm that allows consistent observation of aggressive behaviours with stable monitoring of their body colouration changes. Using this behavioural paradigm, we investigated the relationship between aggressive behaviours and body colouration changes.

**Figure 1.**
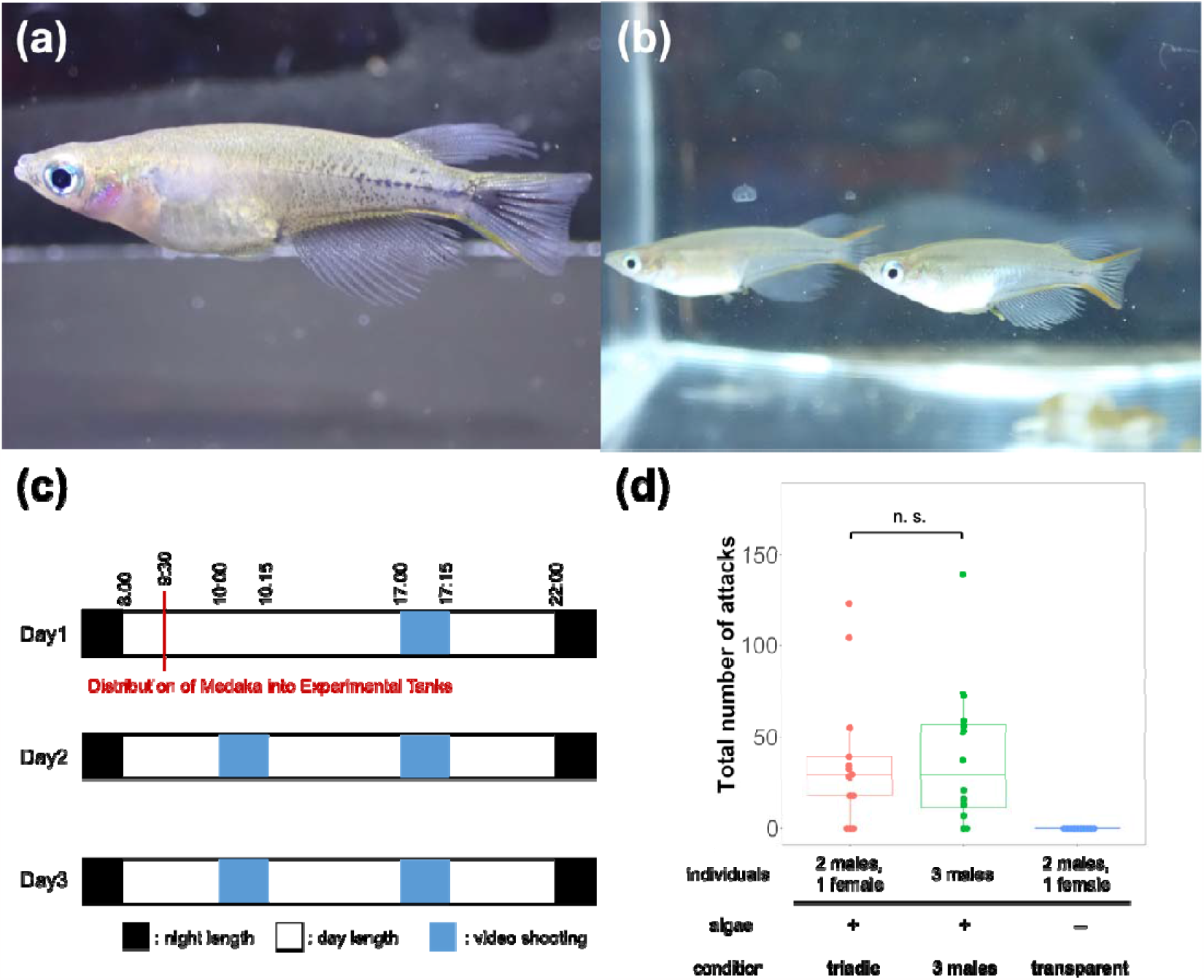
Attacks and body colouration changes in *Oryzias celebensis*. Representative images of male *O. celebensis* with blackened markings (a) and without blackened markings (b). (c) Timetable for the triadic behavioural assays. (d) Number of attacks under the different experimental conditions; algae-covered tank with 2 males and 1 female (triadic), algae-covered tank with 3 males (3-males), and transparent tank with 2 males and 1 female (transparent). Attacks occurred irrespective of the presence of females in the algae-covered tanks, and did not occur at all in the transparent tank under these experimental conditions. \*\**p* < 0.01 and not significant (n.s.) according to the Mann-Whitney *U* test.

## Results

To investigate the experimental conditions that can affect attacks and body colouration changes in *O. celebensis*, we examined the numbers of attacks and the patterns of body colouration changes in small tanks under the following 3 conditions: an algae-covered tank containing 2 males and 1 female (triadic); an algae-covered tank containing 3 males (3-males); and a transparent tank with no algae containing 2 males and 1 female (transparent). The number of attacks in each trial did not significantly differ between the triadic (n = 13) and 3-males (n = 12) conditions (Mann-Whitney *U* test: *Z* = -0.22, *p* = 0.84) (figure 1e), indicating that the fish attack other fish irrespective of the presence of females. In contrast, neither attack behaviour nor black colouration changes were observed in the transparent condition (n = 10) (figure 1b), indicating that the algae-covered walls of the tank were required for the emergence of both the attacks and the black colouration changes.

To investigate whether body colouration correlates with attack frequency, we recorded the number of attacks and the associated body colouration during these attack events for each individual under the triadic condition. The number of attacks by males with blackened markings was higher than that by males without blackened markings or females [generalized linear mixed model (GLMM) followed by Tukey’s post hoc test: black(+)–black(-), estimate = 2.49, SE = 0.952, *p* = 0.0244; black(+)–female, estimate = 4.89, SE = 1.122, *p* < 0.0001; black(-)–female, estimate = 2.40, SE = 1.056, *p* = 0.0598] (figure 2a). We found similar tendencies in the 3-males condition [GLMM followed by Tukey’s post hoc test: black(+)–black(-), estimate = 4.88, SE = 1.01, *p* < 0.0001] (figure 2b). These findings indicate that the *O. celebensis* males with blackened markings exhibited higher aggression toward different conspecific individuals. To determine whether the susceptibility to attacks varies with body colouration, we recorded the number of attacks each individual received, as well as their body colouration at the time of the attack in the triadic condition. The number of attacks received did not differ significantly in relation to body colouration [GLMM followed by Tukey’s post hoc test: black(+)–black(-), estimate = 0.378, SE = 0.574, *p* = 0.788; black(+)–female, estimate = 0.0811, SE = 0.473, *p* = 0.984; black(-)–female, estimate = -0.296, SE = 0.474, *p* = 0.806] (figure 2c). We also found no significant difference between the males with and without blackened markings in the number of attacks received under the 3-males condition [GLMM followed by Tukey’s post hoc test: black(+)–black(-), estimate = 0.132, SE = 0.361, *p* = 0.715] (figure 2d).

**Figure 2.**
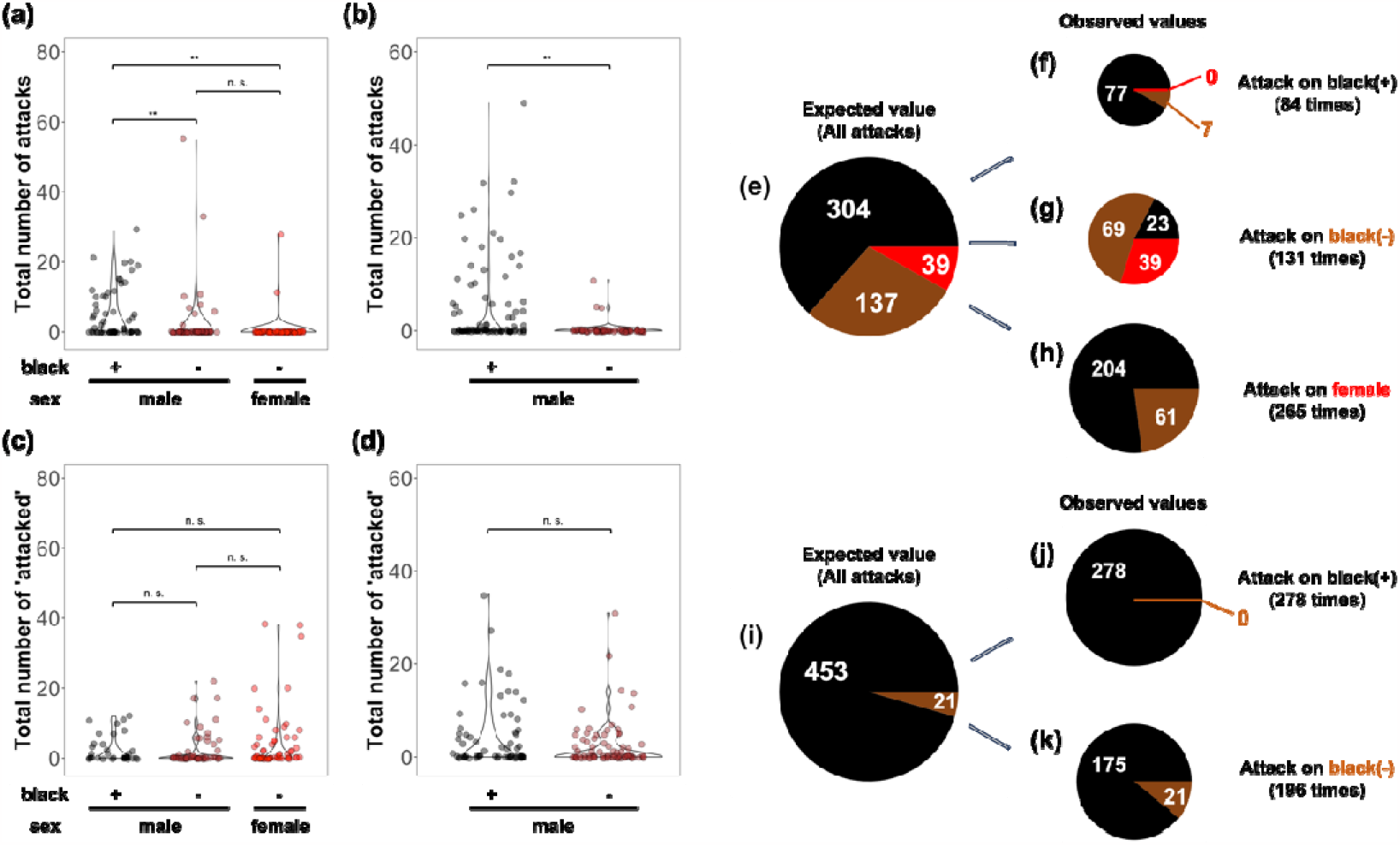
Relationship between attacks and male body colouration. (a, b) Number of attacks under the triadic (2 males and a female) (a) and 3-males (b) conditions. Number of attacks by males with blackened markings was much higher than that by males without blackened markings and females. (c, b) Number of attacks received under the triadic (c) and 3-males (d) conditions. The number of attacks received did not differ significantly between conditions. \**p* < 0.05, \*\**p* < 0.01, and not significant (n.s.) according to generalized linear mixed models followed by Tukey’s post hoc test. (e-k) Pie charts showing directions of attack events under the triadic and 3-males conditions. The left pie chart represents the total number of attacks. The 3 pie charts on the right represent the division of the left pie chart based on the direction of the attacks. The black, brown, and red slices represent males with blackened markings, males without blackened markings, and females, respectively. The size of the slice in the pie charts reflects the number of attacks received.

Next, to examine whether there are biases in the body colouration of the individuals attacked, we analysed the directions of the attack events. In the case of the triadic condition, the observed attack values on males with blackened markings (figure 2f), males without blackened markings (figure 2g), and females (figure 2h) differed significantly from the expected values (figure 2e) [chi-square test: attacks on males with blackened markings: χ^*2*^_2, 84_ = 29.491, *p* < 0.0001; attacks on males without blackened markings: χ^*2*^_2, 131_ = 145.61, *p* < 0.0001; attacks on females: χ^*2*^_1, 265_ = 8.0128, *p* = 0.004645]. In the 3-males condition, the observed attack values on males with blackened markings (figure 2j) and males without blackened markings (figure 2k) also differed significantly different from the expected value (figure 2i) [chi-square test: attacks on males with blackened markings: χ^*2*^_1, 278_ = 12.887, *p* = 0.0003308; attacks on males without blackened markings: χ^*2*^_1, 196_ = 18.279, *p* < 0.0001]. These findings revealed that males with blackened markings were predominantly targeted by other males with blackened markings, while attacks from males without blackened markings or females were rare. On the other hand, males without blackened markings were attacked not only by males with blackened markings but also by other males without blackened markings and females. Additionally, females under the triadic condition experienced a similar attack frequency as males.

## Discussion

The findings of the present study demonstrated that male *O. celebensi*s with blackened body markings exhibited increased aggression toward other members of the same species. In most animals, limited resources such as food, territory, and mates can drive intraspecific competition for their access [9]. The fact that females were often targeted in the triadic condition suggests that the aggressive behavior noted in the present study stems more from competition over resources such as food, rather than from male-to-male competition for mates [10]. In these intraspecific competitions, non-contact aggressive displays may eliminate escalation to physical contact [11, 12]. In some animals, visual threat signals that are specific patterns of behaviors indicate aggressive motivation to assist in resolving conflicts [13-15]. Our findings suggest that the blackened markings on *O. celebensis* males may function as visual cues to signal dominance and fighting ability, thereby aiding in the resolution of disputes over resources.

The link between changes in body colouration and behaviour has been explored in various species. For example, distinct colouration patterns and behavioural displays in the cichlid fish (*Astatotilapia burtoni*) act as visual signals reflecting their state of aggression, which in turn can inhibit the actions of other conspecifics [16, 17, 18]. These changes in colouration and behavior, however, typically take from a few minutes to a day to manifest, possibly due to hormonal influences [19-20]. In contrast, rapid colouration changes occurring within seconds, as seen in chameleons, octopuses, and wrasses, act as immediate, short-term signals of aggression [21-23]. Considering that the body colouration of *O. celebensis* can change within a minute, this species may possess a neural mechanism for exhibiting these visual social signals. In teleost fish, neurotransmitters such as noradrenaline and adenosine control the colouration changes of melanophores [24, 25], suggesting a potential peripheral system for displaying the visual signals. *O. celebensis* could be a promising candidate for applying genome editing techniques, as demonstrated in Japanese medaka (*O. latipes)* [26, 27]. With the availability of a reference genome assembly of *O. celebensis* [8], this species offers a new avenue for probing the molecular and neural mechanisms behind intraspecific communication through rapid colouration changes using advanced molecular and genetic methods such as optogenetics.

## Material and Methods

### Fish and housing conditions

*Oryzias celebensis* (the Ujung pandang strain) was provided by the National Bioresource Project (NBRP) medaka (RS278; https://shigen.nig.ac.jp/medaka/). Fish were maintained in groups in a glass tank (60 cm x 30 cm x 36 cm [height]) containing approximately 30 to 40 individuals with a roughly 1:1 male-to-female ratio, and fed nauplii of brine shrimp or powdered food once a day between 12:00 pm and 1:00 pm. All fish were hatched and bred in our laboratory. Sexually matured male and female medaka 3–15 months of age were subjected to behavioural trials. The water temperature was ∼29 °C and light was provided by LED lights for 14 h per day (08:00–22:00).

### Behavioural trials

We observed aggressive behaviours among 3 adult fish in an acrylic tank (24 cm × 14 cm × 15 cm) for 3 consecutive days (figure 1c) under the following 3 conditions: the triadic condition consisting of 2 males and 1 female in each tank covered with algae on the wall; the 3-males condition consisting of 3 males in each tank covered in algae on the wall; the transparent condition consisting of 2 males and 1 female in each transparent tank without algae on the wall. We used adult fish housed for 2 or more days under the conditions described above. A video of each behavioural trial was recorded twice a day at 10:00 (morning) and 17:00 (evening) using a digital camera (Go Pro Hero9 Black) (figure 1c). On the first morning during the assay (between 9:30 and 10:00), test fish were randomly transferred from the aquarium to each experimental tank. Fifteen minutes after starting the video recording, the test fish were transferred to a small bag with a zip-lock closure, and then photos of the whole body of the test fish were taken using a digital camera (TG-6, Olympus) for individual identification by their fin shapes and pigmentation patterns of the body surface. After obtaining the photo, each fish was returned to their experimental tank. From the first 15 minutes of the video recordings, we quantified aggressive behaviours that were defined as a rapid swim toward a target and the target fleeing as previously described in Japanese medaka (*O. latipes*) [28, 29]. We also described the direction and timing of each attack event, and then determined the body colouration of each test fish while it attacked or was attacked by the other fish. In this study, a male whose markings blackened at least once in each video recording was considered as a male with blackened markings.

### Statistical analysis

We performed a Mann-Whitney *U* test using the “wilcox_test” function in the *coin* package version 1.4.3 implemented in R version 4.2.3 for comparisons of the number of attacks per trial across conditions [30]. Also, to examine whether attack frequencies and attack susceptibility vary based on body colouration, we compared the body colouration with the number of attacks performed and the number of attacks received using R version 4.2.3 with generalized linear mixed models by negative binominal distributions with a log link function using the “glmmTMB” function in the package *glmmTMB* version 1.1.7 [31]. Body colouration of each individual while attacking on while being attacked by other individuals was included as a fixed factor, and experiment tank numbers were included as random factors. For a post hoc test, *P* values adjusted with Tukey’s method were calculated using the package *emmeans* version 1.8.5.

To investigate the possible biases in the body colouration of the individuals that were attacked, we aggregated the number of attacks for each body colouration and analysed the directions of attack events. We set the total ratio of the total number of attacks as an expected value, and the ratio of the total number of attacks categorized by the directions of attack events as observed values. Significant differences between the observed values and expected ones were analysed using the chi-square test implemented in R version 4.2.3.

## Ethics

The work in this paper was conducted using protocols specifically approved by the Animal Care and Use Committee of Tohoku University (Permit Number: 2022LsA-003). All efforts were made to minimize animal suffering, following the NIH Guide for the Care and Use of Laboratory Animals.

## Authors’ contributions

Conceived and designed the experiments: RU, SA, HT. Performed the experiments: RU. Analysed the data: RU, SA, HT. Contributed reagents/materials/analysis tools: SA, HT. Wrote the paper: RU, SA, HT. All authors gave final approval for publication.

## Acknowledgements

We are grateful to the National Bio-Resource Project (NBRP) Medaka for providing the Ujung pandang strain of *O. celebensis* (RS278). We thank N. Sawada for fish care and T. Seki, M. Daimon and D. Kayo for providing advice.

